# Modulating proton sensing by the adenosine receptor A2

**DOI:** 10.64898/2026.05.26.727808

**Authors:** Kyutae D. Lee, Sam Taylor, Daniel G. Isom

## Abstract

For years, it was believed that G-protein-coupled receptors (GPCRs) activated or modulated by changes in physiological pH were mechanistically regulated by pH-sensitive histidine residues on the receptor’s exterior. More recent studies have shown that traditional acid-sensing GPCRs (GPR4, GPR65, GPR68) contain buried acidic amino acids in their 7-transmembrane regions and can be activated by pH-dependent mechanisms. The focus of our research has been the adenosine A2A receptor (A2AR) and its ability to activate at low pH levels without having the typical acidic triad structure found in previously studied acid-sensing GPCRs. We used a combination of bioinformatics and a humanized yeast-based platform, DCyFIR, to identify potential structures involved in the proton-sensing mechanism of A2AR. We also validated some of our A2AR variant yeast phenotypes using mammalian cells. Our data suggest that certain mutations eliminate the pH sensitivity of the A2AR while preserving agonist-induced signaling. Additionally, we demonstrated that one of our mutations (N284D) eliminates pH sensing but increases agonist potency, thereby providing a structural explanation for how pH-sensing and sodium binding occur that differs from previously established models. Overall, these data indicate that G-protein-coupled receptors can sense pH in several ways and demonstrate the potential of a novel, pH-insensitive A2AR in acid-related contexts.

## Introduction

Cells, tissues, and organisms operate within a narrow physiological pH range (pH 5-8) that is essential for life. Proteins detect physiological pH changes through ionizable amino acid side chains. Only five residues are titratable in this range, aspartate, glutamate, histidine, lysine, and arginine, and in solution their pK_a_ values fall well outside the physiological window, except histidine (pK_a_ ≈ 6.6). For this reason, histidine has historically been considered the principal pH-sensing residue, and many proposed mechanisms of acid-activated signaling have centered on extracellular histidine clusters. We have argued elsewhere, and the analysis we present here reinforces, that this view is incomplete^1-3^. When acidic or basic residues are buried in the protein interior, their pK_a_ values can shift by 2 to 4 units in either direction, placing aspartate and glutamate side chains squarely within the physiological pH range^1^. Such residues are rare, conspicuous in three-dimensional structure, and energetically consequential. Each unit of pK_a_ shift corresponds to 1.36 kcal/mol of free energy that can be harnessed to drive conformational change^3,4^. Because their pK_a_ values can be tuned by the local protein environment, buried aspartates and glutamates are, in many respects, better pH sensors than histidines, and the algorithmic identification of such residues in our informatics program, pHinder, has been a productive entry point for finding pH-sensing proteins^1,4,5^.

Our prior work established that GPR4, GPR65, and GPR68 sense protons through a shared buried acidic triad^1^. This triad consists of two structural elements: the DyaD site, a pair of closely spaced aspartates contributed by transmembrane helices 2 and 7 (positions D2.50 and D7.49 in generic Ballesteros-Weinstein [BW] GPCR numbering), and the apEx site, a single buried glutamate on transmembrane helix 4 (E4.53). At physiological pH, the triad is partially protonated, with pK_a_ values shifted to approximately 6-7 by the buried environment. Acid-induced protonation neutralizes the network and stabilizes the active receptor conformation; isosteric mutation of the triad residues to their neutral mimics (aspartate to asparagine, glutamate to glutamine) recapitulates this protonated state and yields pH-insensitive variants that retain signaling. The DyaD site is further bound by Na^+^, which acts as a negative allosteric modulator: sodium binding depresses the pK_a_ values of the paired aspartates and counters acid activation, such that all three receptors function as coincidence detectors of H^+^ and Na^+^. Importantly, the buried acidic triad, defined by the simultaneous presence of D2.50, D7.49, and E4.53, does not occur in any other human GPCR. However, our prior work found that the adenosine receptor 2A (ADORA2A or A2AR), which also lacks the triad, responded similarly to pH as GPR68 in our Dynamic Cyan Induction by Functional Integrated Receptors (DCyFIR) profiling^2,6^. This raised the obvious question motivating the present study: why does A2AR, which lacks the triad, behave so similarly to GPR68, with a pH_50_ in the same physiological range?

Here, we report the molecular basis of acid-activated signaling by A2AR. Combining structural informatics with deep variant profiling in the DCyFIR humanized yeast platform and validation in HEK293T cells, we show that A2AR senses protons through a set of ionizable residues, distinct from the canonical buried acidic triad of GPR4, GPR65, and GPR68, and retains agonist activity for its cognate ligand. Taken together, our results establish that proton sensing in A2AR operates through a non-triad mechanism, demonstrating that GPCRs have evolved multiple structural solutions to detect physiological pH changes. The pH-insensitive A2AR variants we report provide tools for assessing the biological importance of A2A acid-sensing in a broad range of physiological and pathological contexts in which extracellular acidification occurs.

## Results

### Structural informatics identifies candidate proton-sensing residues in A2AR

Buried acidic residues frequently function as effective pH sensors because hydrophobic burial can substantially shift their pKa values, enabling protonation-dependent changes in electrostatic free energy to drive conformational transitions that couple pH to protein structure and function. In GPCRs, this principle is exemplified by GPR4, GPR65, and GPR68, which detect extracellular acidification via a conserved, buried acidic triad mechanism^1^. Notably, this electrostatic architecture appears largely restricted to this subset of proton-sensing GPCRs. We recently found that A2AR also exhibits pH-sensitive signaling behavior despite lacking an analogous buried acidic triad architecture^2^. These observations raised the possibility that A2AR senses pH through an alternative electrostatic mechanism. To investigate this possibility, we used pHinder-based structure-based calculations to identify candidate ionizable residues and electrostatic networks that mediate pH-sensitive behavior within the receptor (Fig. 1).

**Figure 1.**
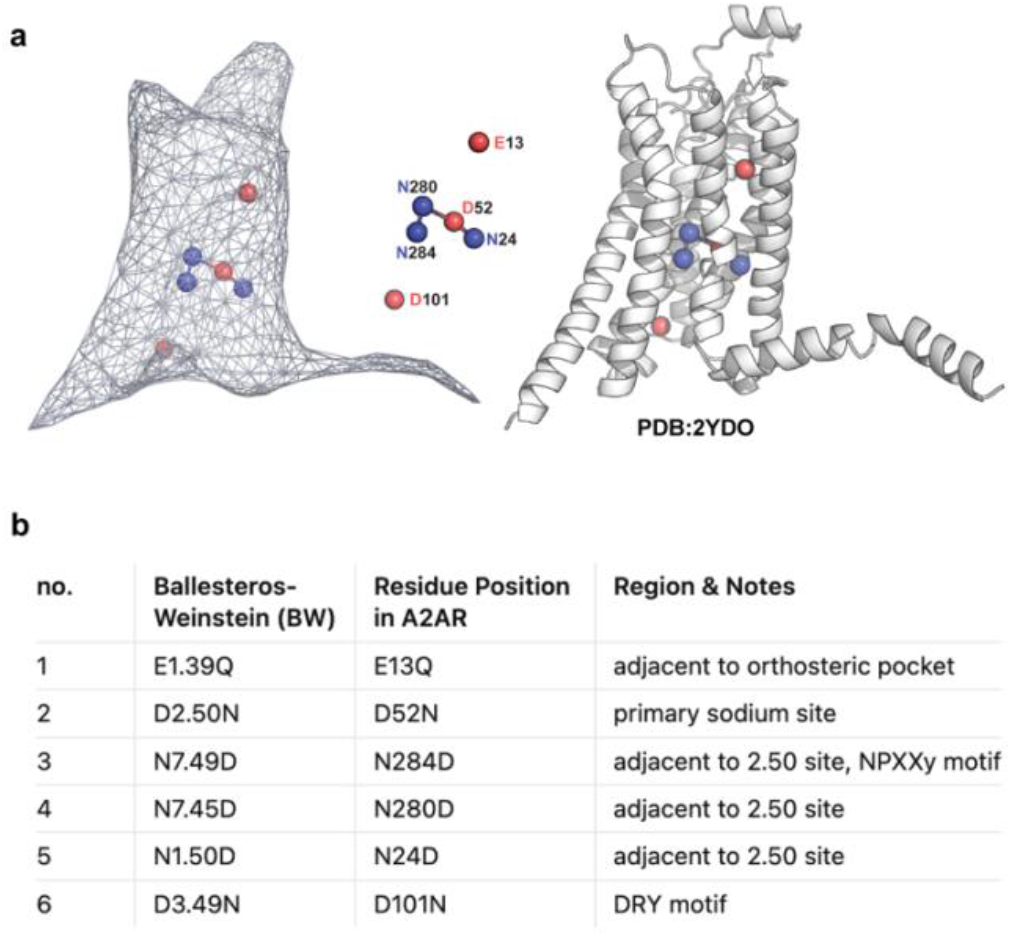
Structural context of A2AR residues targeted for mutagenesis. (a) Spatial arrangement of the six mutated residues within the A2AR transmembrane bundle (PDB: 2YDO). The convex hull surface of the receptor (left) is shown alongside the corresponding cartoon representation (right), with acidic residues (E13, D52, D101) in red and asparagine residues (N24, N280, N284) in blue. D52, N24, N280, and N284 cluster in the canonical sodium-binding pocket near the middle of the helical bundle, while E13 sits near the extracellular orthosteric pocket, and D101 is positioned at the intracellular DRY motif. (b) Summary of the targeted residues, listed by Ballesteros–Weinstein number, position in the A2AR sequence, and structural context.

Application of pHinder to A2AR identified three candidate acidic residues with properties consistent with potential electrostatic sensing roles: E13, D52, and D101 (Fig. 1a, red spheres). Among these, D52 was of particular interest because it participates in sodium-dependent negative allosteric modulation (NAM) and contributes directly to agonist binding and receptor activation. Prior mutational studies have shown that disruption of D52 substantially impairs receptor function. D101 was also notable because it forms part of the highly conserved DRY signaling motif, in which mutations are well known to alter receptor activation and constitutive signaling. Given the established functional sensitivity of D52 and D101, we sought an alternative strategy to modulate receptor electrostatics indirectly rather than directly perturbing these highly conserved acidic residues.

Rather than focusing exclusively on buried acidic groups, we next used pHinder to identify local networks of polar and ionizable residues surrounding D52. These calculations revealed a triad of asparagine residues, N24, N280, and N284, that structurally encase D52 within the receptor core (Fig. 1a, blue spheres). We reasoned that these positions could provide a means to tune the electrostatic environment of D52 through conservative Asn-to-Asp substitutions, in which replacing a single side-chain atom largely preserves steric architecture while introducing a negatively charged carboxylate group. Structural analysis suggested that N24 and N280 form close-range interactions with D52 and would therefore be expected to strongly perturb its electrostatic behavior, potentially disrupting agonist binding and sodium-dependent allosteric regulation. In contrast, N284 is positioned farther from D52 and was predicted to provide a weaker medium-range Coulombic interaction capable of more subtly modulating the local electrostatic environment.

To experimentally evaluate these predictions, we designed a series of rational point mutations intended to minimally perturb steric architecture while altering local electrostatic properties (Fig. 1b). Asp-to-Asn and Glu-to-Gln substitutions were used to mimic protonated acidic side chains by replacing a single oxygen atom while preserving overall side-chain size and hydrogen-bonding geometry: E13Q (adjacent to the orthosteric pocket), D52N (primary sodium site), and D101N (DRY motif). Conversely, Asn-to-Asp substitutions were used to introduce negatively charged character through similarly minimal atomic modification: N24D (N1.50), N280D (N7.45), and N284D (N7.49, within the NPxxY motif). We also generated the N284D/D52N double mutant to test the interdependence of the two strongest candidate sites, as some studies have suggested that N284D rescues D52 mutations^7^. Mutant receptors were evaluated using our previously established yeast-based GPCR signaling system (DCyFIR), which enables quantitative functional characterization of receptor activity in a genetically tractable and experimentally scalable context. Using this approach, we systematically examined how local electrostatic perturbations surrounding D52 influence receptor signaling and pH responsiveness.

### pH sensing is selectively ablated in asparagine-to-aspartate mutants

We screened the mutant library in yeast rather than in mammalian cells for an experimental reason that is central to the interpretation of every pH measurement in this study. In S. cerevisiae, intracellular pH remains essentially invariant across an extracellular pH range of 5.0 to 8.0. In contrast, the cytosol of HEK293T cells acidifies by approximately one pH unit over the same extracellular range. This invariance is decisive for our experiments, because the protonation states of ionizable residues elsewhere in the cell, including those of downstream signaling components (e.g., G-proteins), would otherwise confound the results. By performing the primary screen in the DCyFIR humanized yeast platform, in which human GPCRs are genetically introduced and coupled to a pheromone-pathway transcriptional reporter (mTq2), we localized the pH dependence of the assay to the extracellular-facing receptor surface and the receptor interior. This is the same advantage that enabled us to dissect the buried acidic triad of GPR4, GPR65, and GPR68 in our previous work^1^.

We first asked which mutants retain pH sensing, that is, the capacity to mount an acid-driven response in the presence of very low adenosine concentration (Fig. 2). Plotting mTq2 fluorescence as a function of extracellular pH across the range 5.0 to 7.0, wild-type A2AR produced a robust acid-activated response, with signal increasing approximately 2.5-fold from pH 7.0 to pH 5.0 (Fig. 2a, b). Three of the seven mutants retained measurable pH responses: D52N and E13Q preserved 71% and 53% of the wild-type acid-induced fold change, respectively (Fig. 2c), while D101N maintained a strongly elevated absolute signal across the entire pH range but showed little additional pH sensitivity, consistent with the constitutive activity expected from a DRY-motif mutation. The four remaining mutants, N24D, N280D, N284D, and the N284D/D52N double mutant, displayed flat, near-background basal profiles, with no detectable pH-driven change in signal. The selective elimination of pH sensing in three of the four asparagine-to-aspartate substitutions confirmed that introducing a carboxylate into the local environment of D52 is sufficient to ablate the receptor’s acid response, validating the structural prediction that the N24/N280/N284 triad participates in A2AR proton sensing.

**Figure 2.**
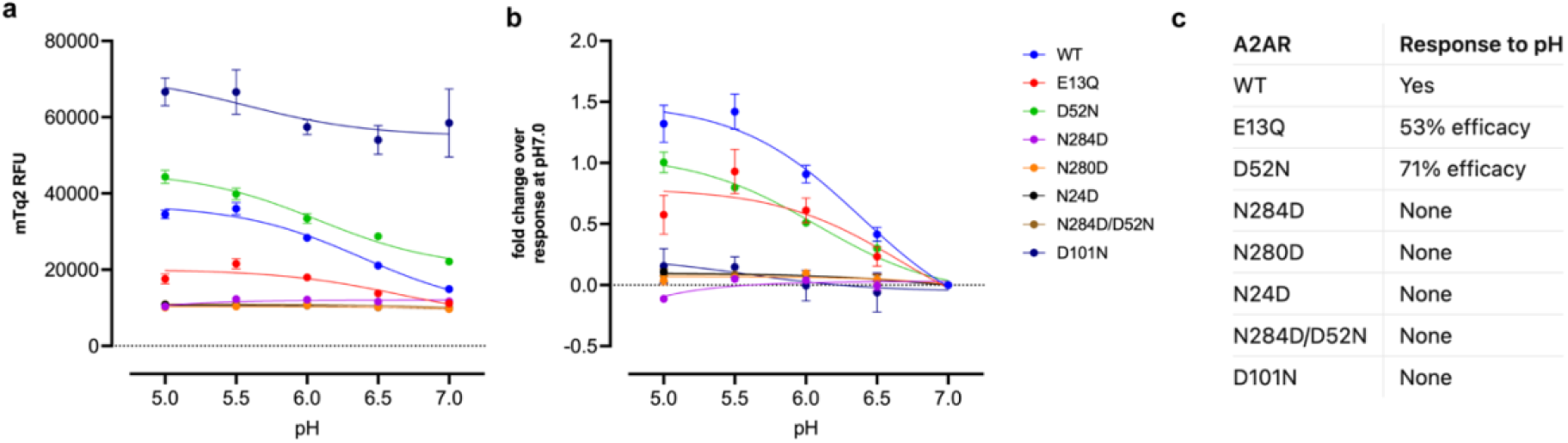
A2AR signaling is differentially sensitive to extracellular pH in the DCyFIR humanized yeast platform. (a) mTq2 relative fluorescence units (RFU) in 10nM of adenosine plotted as a function of extracellular pH for wild-type (WT) A2AR and seven-point mutants expressed in the Dynamic Cyan Induction by Functional Integrated Receptors (DCyFIR) humanized yeast platform, in which mTq2 serves as a transcriptional reporter of human GPCR signaling. (b) The same data were normalized as fold change relative to the response at pH 7.0, isolating the pH-driven component of each variant’s signaling activity. (c) Summary classification of each variant’s pH response. Data represent mean ± SD from n=4 independent biological replicates.

### Dose–response profiling across pH separates loss of pH sensing from loss of receptor function

The flat pH profiles observed in Fig. 2 could, in principle, reflect either selective loss of pH-sensing or generalized receptor dysfunction. To distinguish these possibilities, we measured full adenosine concentration–response curves for all eight receptor variants at each of five extracellular pH values (pH 5.0, 5.5, 6.0, 6.5, and 7.0; Fig. 3). This revealed which mutants are signaling-competent and asked, at every pH, what their adenosine-stimulated response looks like.

**Figure 3.**
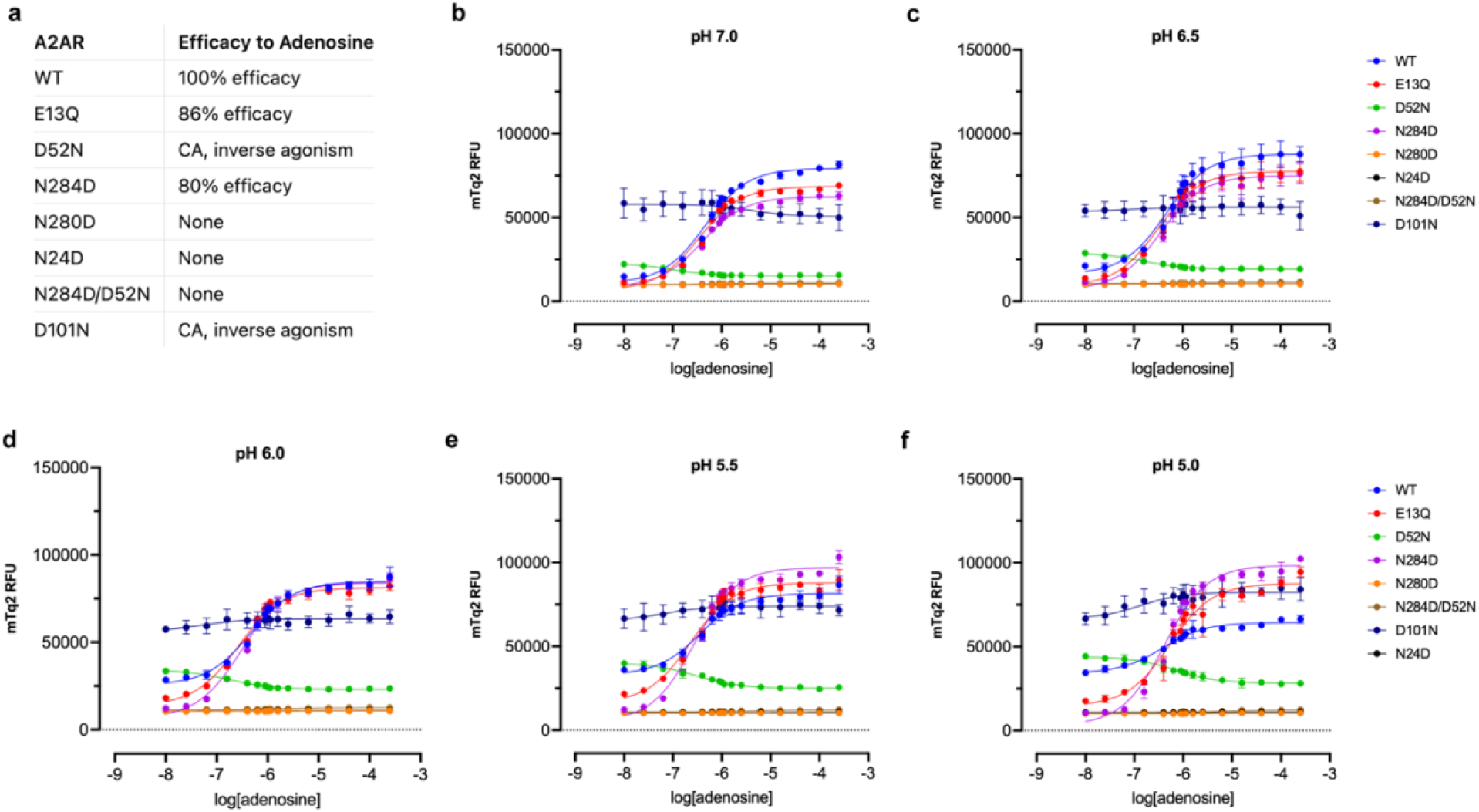
A panel of A2AR mutants targeting conserved acidic and polar residues reveals distinct adenosine response profiles across physiological pH. (a) Summary of adenosine efficacy for wild-type A2AR and seven-point mutants. (b–f) Adenosine concentration–response curves for all eight receptor variants measured at (b) pH 7.0, (c) pH 6.5, (d) pH 6.0, (e) pH 5.5, and (f) pH 5.0 in the DCyFIR humanized yeast platform, using mTq2 as a transcriptional reporter of A2AR signaling. Signals are plotted as raw mTq2 relative fluorescence units (RFU). Data represent mean ± SD from n=4 independent biological replicates.

Four distinct phenotypes emerged (Fig. 3a). Wild-type and E13Q produced full agonist responses at all pH values (WT: 100%, E13Q: 86% efficacy relative to WT at pH 7.0). N284D produced an 80% response at pH 7.0 and was notable for showing a maximal response that increased rather than decreased at acidic pH, the opposite of wild-type behavior. D52N and D101N showed elevated basal signals that decreased upon adenosine addition, a hallmark of constitutive activity coupled with inverse agonism by adenosine, and this behavior became more pronounced at lower pH. The remaining three variants (N24D, N280D, and the N284D/D52N double mutant) produced flat, near-baseline responses at all pH and adenosine concentrations tested, indicating loss of function.

N284D and E13Q retain robust adenosine-stimulated signaling but have lost the wild-type pH dependence of that signaling, providing the first A2AR variants in which proton sensing is selectively decoupled from receptor activation. N24D, N280D, and N284D/D52N are signaling-incompetent and therefore uninformative for the pH question. Still, their loss of function is internally consistent with the structural information in that close-range carboxylate substitution around D52 (N24, N280) should strongly disrupt receptor function. In contrast, medium-range, outward-facing substitution (N284) should be more permissive. These results suggest that we effectively removed A2AR’s pH-sensing capabilities while minimally altering its agonist-induced signaling activity.

### Mutant-specific reshaping of the pH–response landscape

To examine the pH dependence of the three signaling-competent informative variants more directly, we replotted their dose–response data as fold change over basal at each pH (Fig. 4a–c) and overlaid the pH 7.0 and pH 5.0 curves normalized to maximal response (Fig. 4d–f). Wild-type A2AR signaling was progressively suppressed by acidification, with maximal response falling roughly threefold between pH 7.0 and pH 5.0 (Fig. 4a). The N284D mutant displayed the opposite behavior: acidic conditions enhanced rather than suppressed the maximal adenosine response, reaching nearly tenfold over basal at pH 5.0 (Fig. 4b). E13Q showed an intermediate, partially attenuated pH dependence, with maximal responses compressed across the pH range relative to wild-type (Fig. 4c).

**Figure 4.**
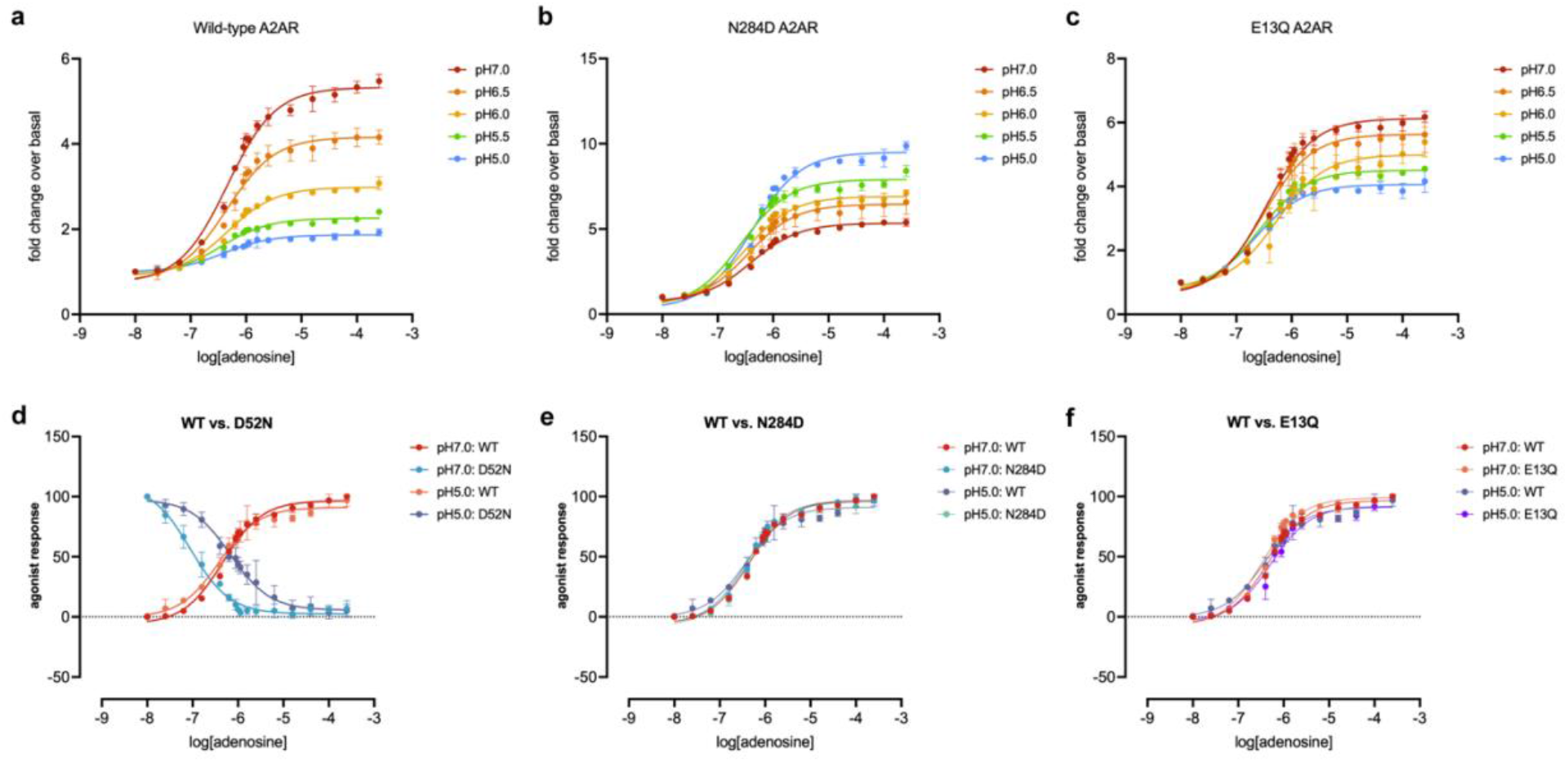
Extracellular pH modulates adenosine-induced A2AR signaling in a mutant-specific manner. (a–c) Adenosine concentration–response curves measured at pH 5.0 to 7.0 normalized as fold change relative to the response at the lowest adenosine dose in the DCyFIR humanized yeast platform expressing (a) wild-type A2AR, (b) the N284D mutant, or (c) the E13Q mutant, with mTq2 reporter signal plotted as fold change over signaling at 10nM. (d–f) Direct comparisons of wild-type and (d) D52N, (e) N284D, and (f) E13Q mutant receptors at pH 7.0 and pH 5.0, normalized to maximal agonist response. These graphs are the same data as in Figure 3, for a clearer depiction of changes in maximal response rather than agonist potency. Data represent mean ± SD from n=4 independent biological replicates.

The normalized overlays in Fig. 4d–f isolate the source of these differences. For D52N (Fig. 4d), the pH 5.0 curve is inverted relative to wild-type, with apparent agonist response decreasing as adenosine is added, consistent with the inverse-agonism phenotype noted above. For N284D (Fig. 4e) and E13Q (Fig. 4f), the pH 7.0 and pH 5.0 normalized curves overlap closely with wild-type, indicating that the pH-dependent differences in panels b and c reflect changes in maximal response (efficacy) rather than in agonist potency. The mutations, therefore, reshape the pH–efficacy relationship of A2AR without altering its apparent affinity for adenosine, a finding consistent with the interpretation that the targeted residues contribute to the energetics of the active-state conformation rather than to ligand binding directly.

### Mammalian-cell validation using a luminescence-based cAMP biosensor

To confirm that the yeast findings translate to a mammalian model, we expressed the most informative mutants in HEK293T cells and measured cAMP accumulation using the GloSensor luminescence-based biosensor (Fig. 5). NECA, a potent synthetic agonist for adenosine receptors, was chosen because adenosine deaminase must be used in the assays, otherwise the cAMP biosensor is saturated from the existing adenosine produced endogenously by the cells. We selected wild-type A2AR together with E13Q, D52N, and N284D, the three variants that produced the clearest and most distinct phenotypes in the yeast screen. NECA dose-response curves confirmed that the three mutants behave as predicted by the yeast data and are indeed functional. As expected, D52N produced no detectable concentration-dependent response across the tested range; its curve overlapped completely with the control, confirming that the residual high-concentration signal in D52N-expressing cells reflects background biosensor activity from endogenous adenosine receptors rather than overexpressed–receptor–mediated cAMP production. Wild-type and E13Q produced overlapping dose-response curves, consistent with their similar adenosine efficacies in yeast. Interestingly, N284D displayed a leftward-shifted curve, indicating increased agonist potency in the mammalian context. These data suggest that raising the local pKa of D52 by the N284D mutation leads to greater protonation/neutralization of D52 and thus decreased negative allosteric modulation by sodium, resulting in increased potency for the N284D mutant.

**Figure 5.**
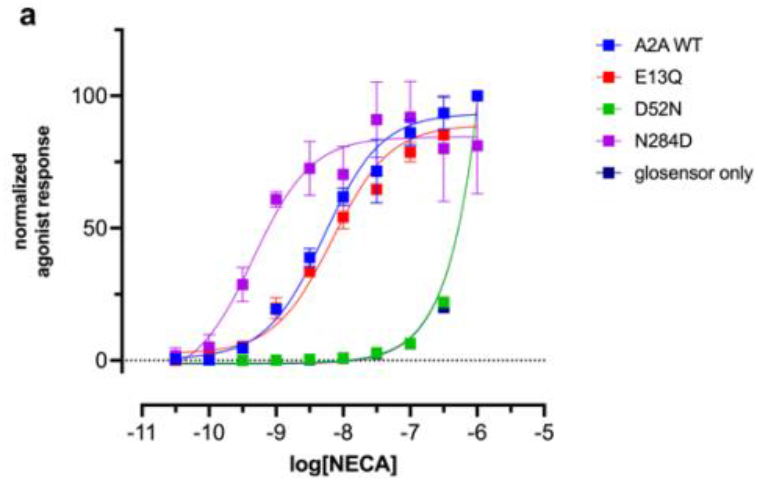
Measuring agonist sensitivity of A2AR mutants using a GloSensor cAMP assay in a mammalian cell system. (a) Concentration–response curves for NECA-induced cAMP accumulation at wild-type A2AR and three A2AR point mutants (E13Q, D52N, N284D) expressed in HEK293T cells alongside the GloSensor 22F cAMP biosensor. Responses are normalized to the maximal wild-type signal. The “glosensor only” control overlaps completely with D52N, confirming that the residual high-concentration signal in D52N-expressing cells reflects background biosensor activity of endogenous adenosine receptors rather than overexpressed receptor-mediated cAMP production. Data represent mean ± SD from a representative experiment of n=2 independent experiments.

These data establish that the agonist-response phenotypes observed in yeast, preserved signaling with altered pH dependence for E13Q and N284D, loss of agonist-induced signaling for D52N, are recapitulated in a mammalian cell system. The convergence between the yeast and HEK293T results supports the broader conclusion that the buried ionizable residues identified by pHinder participate in A2AR pH sensing through a mechanism that is structurally distinct from, but operationally analogous to, the buried acidic triad of GPR4, GPR65, and GPR68.

## Discussion

The principal conclusion of this work is that GPCRs have evolved multiple molecular solutions to the problem of detecting physiological pH. GPR4, GPR65, and GPR68 sense protons through a shared buried acidic triad, paired DyaD aspartates at D2.50 and D7.49, and an apEx glutamate at E4.53, which is absent from every other human GPCR we have analyzed, including A2AR^1^. The triad is allosterically tuned by sodium binding and behaves as a coincidence detector of H^+^ and Na^+1^. A2AR, in contrast, behaves as an acid-activated GPCR with a pH_50_ comparable to GPR68 despite lacking the triad architecture entirely^2^. The mutational dissection we present here identifies a set of buried and conformationally constrained ionizable residues, most prominently N284 on transmembrane (helix 7), in which a change to its acidic counterpart selectively eliminates pH sensing without disrupting agonist-stimulated signaling. The mechanism that emerges is structurally distinct from the triad and cannot be summarized as a single ionizable network, as in GPR4, GPR65, and GPR68. We anticipate that as more GPCRs are characterized at this level of detail, additional acid-activated receptors will be found to operate through still other mechanisms, and that the classical extracellular-histidine model, which has dominated the field for nearly two decades, will be revealed as one minor contributor among several.

The pH-insensitive, but agonism-retaining A2AR variants we report (N284D, E13Q), provide an experimental handle for exploring the biological role of A2A acid sensing. A2AR is one of the most extensively characterized receptors in pharmacology^8^, with established functions in cardiovascular regulation^9^, neurotransmission^10^, immune cell suppression in the tumor microenvironment^11^, and protection against ischemic injury^12^. Each of these contexts involves substantial local acidification: tumor microenvironments routinely fall below pH 6.5, ischemic tissue acidifies within minutes, and the endolysosomal pathway through which internalized receptors signal can drop to pH 4.5 - 5.0 in mature lysosomes^13^. Until now, it has not been possible to ask whether A2AR signaling under these conditions depends on the receptor’s intrinsic acid sensitivity or whether it is driven exclusively by adenosine concentration, because no separable pH-insensitive variant existed. Our variants now make this distinction experimentally accessible. In vivo comparisons between wild-type and N284D A2AR in tumor-bearing animals, in ischemia–reperfusion models, and in conditional cell-type-specific systems should resolve whether the receptor’s acid sensitivity contributes to immune suppression in the tumor microenvironment or to cardioprotection during ischemia. In vitro, the same variants enable scalable cell-based screening for adenosine-pathway compounds without the confounding effects of pH on receptor activity, a practical advantage for drug discovery efforts that target A2A in pathological tissues where pH is itself dysregulated.

Several of these positions have prior functional precedent in the GPCR mutagenesis literature, which informed our expectations for each variant. At D2.50 (D52 in A2AR), alanine substitution abolishes G protein–dependent signaling at A2AR^14^, and neutral substitution at D2.50 disrupts or impairs signaling across diverse class A GPCRs (reviewed in Katritch et al., 2014); the Asn substitution we used here preserves the side-chain volume and hydrogen-bonding geometry while removing the negative charge. At N7.49 (N284 in A2AR), alanine substitution similarly abolishes A2AR signaling^14^, and an Asp-to-Asn mutation at D2.50 could be partially rescued by a compensating Asn-to-Asp substitution at N7.49 in the 5-HT2A receptor^7^, indicating that the two positions function as an interacting electrostatic pair. At N7.45 (N280 in A2AR), alanine substitution at A2AR reduces agonist-stimulated signaling, with a non-significant trend toward elevated basal activity^14^. At D3.49 (D101 in A2AR), neutral substitution of the conserved DRY-motif acidic residue produces constitutive activity across multiple class A GPCRs, including rhodopsin (E3.49Q^15^), the β2-adrenergic receptor (D3.49N^16^), and the α1B-adrenergic receptor (D3.49A^17^). To our knowledge, mutations at E1.39 (E13 in A2AR) and N1.50 (N24 in A2AR) have not been systematically characterized with respect to receptor function.

The structural location of the residues most strongly implicated in A2AR pH sensing is worth considering in its own right. N284 sits on transmembrane helix 7 within the conserved NPxxY motif (N7.49), immediately adjacent to the cytoplasmic activation switch of class A GPCRs. The fact that an N-to-D substitution at this site abolishes pH sensing while simultaneously increasing agonist potency is consistent with a model in which N284 contributes to the energetics of the inactive-to-active conformational transition, and in which protonation of nearby acidic residues at low pH normally biases the receptor toward the active state. The clustering of pH-sensitive asparagines around the primary sodium-binding pocket, N24 on transmembrane helix 1 (N1.50) and N280 on transmembrane helix 7 (N7.45), both in close-range contact with D52 (D2.50), raises the possibility that the pH-sensing network is centered on the canonical sodium-binding site itself, with D52 acting as the electrostatic linchpin and the surrounding asparagines tuning its protonation behavior. This is mechanistically distinct from the triad architecture, in which the DyaD and apEx sites are spatially separated and activate through a distinct conformational route. The fact that the N284D/D52N double mutant is completely signaling incompetent, whereas either single mutation alone retains at least some signaling, is consistent with these residues acting through a shared network rather than as independent contributors. Whether the A2AR network is best described as a single distributed pH sensor or as a set of coupled ionizable contributions whose effects sum at the level of receptor activation remains to be determined; high-resolution structural studies of the pH-insensitive variants in active and inactive states, together with molecular dynamics calculations parameterized for the relevant pK_a_ values, should resolve this question.

Sequence analysis across class A GPCRs lends independent support to this interpretation. Position 7.49 is conserved as asparagine in roughly 86% of class A receptors, but approximately 7%, including PAR1 and 51 other receptors, carry an aspartate at both 2.50 and 7.49^18^. The evolutionary tolerance of an acidic residue at 7.49 in this subset of receptors, together with the demonstration that a D2.50N/N7.49D double mutant can rescue function in 5-HT2A^7^ and μ-opioid^19^ receptors, indicates that the 7.49 position is fundamentally compatible with a carboxylate side chain. However, our finding that the equivalent A2AR double mutant (D52N/N284D) is entirely signaling-incompetent stands in stark contrast to the functional rescue observed in the 5-HT2A and μ-opioid receptors. We propose that this divergence reflects A2AR’s specialized role as an agonist-independent pH sensor. In A2AR, the basal conformational state is poised on a finely balanced electrostatic edge, relying on the precisely tuned pKa of D52 within its specific asparagine microenvironment. While swapping the acidic residue to position 7.49 preserves the net charge required for general class A GPCR activation, it disrupts the highly specific spatial geometry required for A2AR proton sensing. Without the mutual electrostatic network formed by both residues, the active-state equilibrium collapses. Thus, the lack of rescue underscores that the A2AR sodium pocket is not merely a structural linchpin, but an evolutionarily specialized microenvironment tuned specifically for proton detection. This evolutionary plasticity is consistent with our finding that introducing an aspartate at N284 abolishes pH sensing while preserving signaling: the position appears to tolerate a negatively charged side chain, and its substitution here selectively reshapes the local electrostatic environment of the sodium-binding pocket without disrupting the structural framework required for activation. Whether the receptors that naturally carry D7.49 also exhibit pH-modulated signaling, and whether N7.49D variants more broadly confer pH-insensitive behavior on other class A GPCRs, are questions our platform is well-suited to address.

Finally, we expect that A2AR is not the last GPCR to be added to the list of proton-activated human receptors, and that the larger picture will look less like a small set of dedicated acid sensors and more like a continuum of proton-modulated receptors. Our recent work on proton-gated coincidence detection showed that more than 100 individual GPCR–Gα coupling combinations are modulated, positively or negatively, by physiological pH changes, thereby affecting agonist potency, efficacy, cooperativity, antagonism, and constitutive activity. A2AR and GPR4/65/68 represent the extreme end of this continuum, in which acid activation occurs in the absence of agonist; many more receptors are likely to function as graded or Boolean-like proton-gated detectors that integrate H^+^ with their canonical ligand. The yeast platform we have developed is well-suited to systematic characterization of this larger landscape, particularly across the non-human GPCR sequence space, where novel pH-sensing architectures may have evolved in response to organism-specific microenvironments. Combining structural informatics with high-throughput functional profiling, as reported here for A2AR, should make it feasible to map the full spectrum of GPCR proton responses and, in doing so, to identify the molecular sensors through which extracellular and endosomal pH signals are transduced into cellular decisions.

## Methods

### Mammalian Cell Culture and Transfection

Human cell lines were obtained from ATCC, collaborators, or existing stocks and cultured under recommended conditions. HEK293T cells were cultured in DMEM (Thermo Fisher Scientific, Cat# 11995065) with 10% FBS (HyClone, #SH30071.03) and 1% Penicillin-Streptomycin-Glutamine (PSG, Cat# 10378016). All cells were incubated at 37 °C with 5% CO_2_ and passaged using TrypLE Express (Thermo Fisher, Cat# 12604013). For transient expression experiments, cells were seeded in 6-well plates and transfected using TransIT (Mirus Bio #MIR 2305). After 24 h, cells were trypsinized, counted, and replated into white 384-well plates (Greiner Bio-One #781095) at 5,000–10,000 cells per well for downstream assays.

### Yeast DCyFIR Strains

The yeast strains used in this study are identical to those previously described^6^. Briefly, we genetically engineered the wild-type yeast strain BY4741 to include 1) deletion of the native yeast GPCR (ste2Δ) and its negative regulators of the pheromone pathway (GTPase activating protein sst2Δ, cell cycle arrest factor far1Δ); 2) installation of a CRISPR addressable expression cassette in chromosome locus X2 that includes a constitutive promoter (PTEF), synthetic universal targeting sequence for CRISPR/Cas9 editing (i.e., knock-in of human GPCRs), and terminator (TCYC1B); and 3) installation of the mTurquoise2 transcriptional reporter in place of the pheromone responsive Fig1 gene (fig1Δ:: mTurquoise2). After genetic modification of this base strain to remove these elements, we expanded the number of GPCR-Gα strains to ten through CRISPR-Cas9 genome engineering to add human C-terminal sequences to the native yeast Gα subunit Gpa1. Finally, we introduced human GPCR genes into the genomes of the ten GPCR-Gα strains using high-throughput CRISPR-Cas9 genome engineering.

### Media

All yeast strains were streaked onto YPD media plates (20 g/L peptone, 10 g/L yeast extract, 2% glucose, 15 g/L agar) from a frozen 30% glycerol stock at −80°C. All DCyFIR screen experiments and pH titration studies utilized yeast that were cultured in low-fluorescence synthetic complete drop-out media (SCD media; 50 mM potassium phosphate dibasic, 50 mM MES hydrate, 5 g/L ammonium sulfate, 1.7 g/L yeast nitrogen base w/o amino acids, folic acid, and riboflavin (Formedium; CYN6505), 0.79 g/L complete amino acid mix (MP Biomedicals; 4500022), and 2% glucose) titrated to the desired pH by adjusting with HCl or NaOH. In addition to the use of SCD media for yeast growth in pHluorin-based studies where intracellular pH of yeast was assessed through measurement of pHluorin fluorescence, yeast were also plated on selective media (SD media lacking leucine (5 g/L ammonium sulfate, 1.7 g/L yeast nitrogen base without amino acids (MP Biomedicals; 4510522), 1 NaOH pellet (VWR; BDH9292), 0.69 g/L CSM-LEU (amino acid mix lacking leucine), 2% glucose, and 15 g/L Bacto Agar for plates)) that had been filter sterilized before seeding. All media, except YPD (autoclaved), were 0.2 μm filter-sterilized.

Phosphate Buffered Saline (PBS) was prepared by creating two individual solutions of mono-and dibasic potassium phosphate (100 mM potassium chloride, 25 mM potassium phosphate). Solutions of different pH values of PBS were prepared individually by mixing the mono- and dibasic solutions until the desired pH was achieved, as measured with an Accumet XL150 pH meter (Fisher Scientific; Hampton, NH).

### Plasmids

The Rajini Rao Lab at Johns Hopkins University provided the yeast plasmid pYEplac181-pHluorin. The pHluorin gene from pYEplac181-pHluorin was cloned into the pLIC-His vector (pLIC-His-pHluorin) so that the pHluorin could be expressed recombinantly and purified from E. coli. The mammalian plasmid pcDNA3.1(+)-pHluorin was generated by GenScript (Piscataway, NJ) by synthesizing and cloning a codon-optimized version of the pHluorin gene into the pcDNA3.1(+) backbone. Human A2AR cDNA from the PRESTO-TANGO library was sub-cloned into the pYEplac181 vector using the NEBuilder HiFi DNA Assembly Kit (New England BioSciences, E2621S) to create a yeast 2µ high-copy plasmid for expressing human A2AR (pYEplac181-A2AR). pcDNA3.1(+)-ADORA2A was “dePresto” by using the previously described method to remove assay-specific protein and tags from the C-terminus^20^.

All mutants were created from the pYEplac181-A2AR and pcDNA3.1(+)-ADORA2A backbones using the strategy called the 5’-phosphate PCR Assembly. Briefly, vectors were amplified using Q5 High-Fidelity DNA Polymerase (NEB) reactions with a 5’-phosphate-modified complementary primer, a primer containing the desired change, and a complementary region (in opposite orientations), and the template plasmid DNA. PCR products were verified by agarose gel electrophoresis, then digested with DpnI (NEB) at 37°C for 1.5 hours, followed by heat inactivation. Ligation was performed overnight at 16°C or for two hours at room temperature using T4 DNA Ligase (NEB). Ligase was heat-inactivated, and 2µL of the ligation mix was transformed into competent DH-5α cells.

All plasmids were verified via Sanger or nanopore sequencing (Eurofins Genomics).

### DCyFIR Profiling

Solutions. 1× TE buffer (10 mM Tris, 1 mM EDTA). Lithium acetate (LiOAc) mix (100 mM LiOAc, 10 mM Tris, 1 mM EDTA). PEG mixture (40% (w/v) PEG 3350, 100 mM LiOAc, 10 mM Tris, and 1 mM EDTA).

Strain Generation. Yeast cells were initially inoculated into 3 mL of YPD medium and incubated overnight at 30 °C with shaking. The following day, 100 µL of this starter culture was transferred into 5 mL of fresh YPD and incubated for approximately 2.5 hours until the culture reached an optical density (OD) of 0.6–1.2. The cells were harvested, and the resulting pellet was washed sequentially with 5 mL of 1× TE buffer and 5 mL of the LiOAc mix. After the final wash, the yeast pellet was resuspended in 200 µL of LiOAc buffer for transformation.

For each transformation reaction, 50 µL of the chemically prepared yeast cells were combined with 300 ng of plasmid DNA, 20 µL of repair DNA or donor payload, 350 µL of PEG mix, and 5 µL of salmon sperm DNA in a sterile 1.5 mL microcentrifuge tube. The samples were briefly vortexed and incubated at room temperature for 30 minutes. Following this incubation, 24 µL of 100% DMSO was added to the mixture, and the cells were subjected to a heat shock at 42 °C for 15 minutes. Post-heat shock, the cells were pelleted by centrifugation at 10,000 rpm for 3 minutes, gently resuspended in 100 µL of YPD without vortexing, and plated onto leucine-depleted selective media for approximately 3 days of growth at 30 °C.

Strain Growth. Before DCyFIR profiling, each A2AR-variant-Gα strain was streaked onto YPD plates and incubated at 30°C for 2 days. One colony was picked from each plate and seeded into 1 mL of SCD media in a 96-well deep-well plate. Cultures were incubated overnight at 30 °C in a static environment. The cells were then diluted down to an OD600 of approximately 0.05 at both pH levels and grown to mid-log phase during the day. The cells were then diluted down again to an OD600 that would give them an OD600 of 1 when harvested 24 hours later. This usually corresponds to a 16-18 hour period of growth under non-shaking conditions at 30 °C. It should be noted that there is a difference in doubling time between pH 7 and pH 5, requiring a greater initial cell concentration at pH 7. Experimental Setup, Measurements, Replicates. Following the production strain growth procedure described above, the cells were collected and adjusted to a uniform concentration based on an OD600 reading of 0.1. These normalized cultures were divided equally among four wells of a three-eighty-well plate as follows: forty-five microliters of each normalized culture per condition was placed within four wells of a three-eighty-well plate to generate four independent replicate sets while generating a total of sixteen data points for each condition: four data points for each vehicle condition at each pH level and four data points for each agonist condition at each pH level. Each well received 40 microliters total after adding 4 microliters of vehicle or a 10× dilution of the agonist. The cultures were allowed to continue growing for an additional eighteen hours at pH five and twenty-four hours at pH seven until they reached an OD600 value ranging from three to four. Because the rate of growth is slightly less at pH seven than at pH five, due to differences in doubling time, it was necessary to continue growing some subsets of the receptor population an additional twenty-four hours beyond the eighteen hours required for cells grown at pH five for them to reach an OD600 value range of three to four. Therefore, all DCyFIR profiling results shown in Figs. 2, 4, and 5 were obtained from cultures matched for OD600. GPCR signaling activity was determined through measurement of fluorescence emitted from the mTq2 transcriptional reporter through quantitative measurement of the fluorescence of the OD-matched cultures utilizing a ClarioStar microplate reader with the following settings: Bottom Read; Ten Flashes/Well; Ex-F: 430 – 10nm; Dichroic Filter: LP458nm; Em-F:482 – 16nm; Gain =1300.

### pH Titration Experiments

Strains were grown. Each GPCR-Gα strain was taken out of a glycerol stock on YPD plates and incubated at 30°C for 48 hours. A single colony for agonist-treated pH titrations was selected and transferred to 5 ml of SCD media at pH 5.0, 5.5, 6.0, 6.5, and 7.0. The same process was followed to select a single colony that would be added to 5 ml of SCD media at pH 5.0 or 7.0 for constitutively active GPCR-Gα pH titrations based upon the level of activation by low or high pH, respectively. Cells grown in each production run were diluted to an optical density at 600 nm of 0.05 (for pH 5.0, 5.5, and 6.0 growths) or 0.1 (for pH 6.5 and 7.0 growths), using each of the different pH media in a 50 ml conical tube, shaken at 30°C until mid-log phase was reached during the day. On the second day, cells were back-diluted so that they will be at an OD600 value of 1.0 after 4-6 hours, with the use of fresh media at the desired time after diluting overnight-shaken cultures at 30°C, so that they are completely adapted to the pH of the media. Then the cultures were subject to DCyFIR profiling methods as mentioned above.

### pHluorin Calibration

A pHluorin standard curve was calculated using purified ratiometric pHluorin. 200 μL PBS buffer titrated to each pH (137 mM NaCl, 2.7 mM KCl, 10 mM Na_2_HPO_4_, 1.8 mM KH_2_PO_4_) supplemented with 20 mM MES (Sigma-Aldrich, #M3671) or 20 mM HEPES (Sigma-Aldrich, #H3375, adjusted to the indicated pH values using HCl or NaOH) were placed into a 96-well microplate (CytoOne; CC7626-7596), and 36 μL aliquots were moved in technical quadruplicate to a black bottom 384-well plate (Greiner, 781096). Briefly, 4 μL ratiometric pHluorin (1 μM) were resuspended in each pH PBS buffer. The plate was vortexed at 2000 rpm for 30 seconds before reading. Excitation spectra were collected using a ClarioStar microplate reader (BMG LabTech) with the following parameters: top read, 40 flashes/ well, excitation start: 350 to 10 nm, excitation end: 495 to 10 nm, 5 nm steps; emission: 520 to 10 nm; instrument gain: 1700; focal height 7 mm. Raw fluorescence values at 385 and 475 nm were used to calculate the pHluorin ratio (385/475 nm) for each technical replicate. A standard curve was built by plotting the pHluorin ratio as a function of pH and fit using a sigmoidal 4PL model (X = log[concentration]) in GraphPad Prism. Experimental error was calculated in GraphPad Prism as the SD of the mean for the n=4 technical replicates.

### Acid Stress Assay and Phenotypic Classification

For acid stress assays, the HEK293T cells was seeded into the aforementioned white Greiner 384-well plates at a density of 5,000–10,000 cells per well and allowed to adhere for 48 h post-transfection or post-transduction. Cells were washed with pre-warmed PBS and incubated in pH-adjusted buffer (137 mM NaCl, 2.7 mM KCl, 10 mM Na_2_HPO_4_, 1.8 mM KH_2_PO_4_) supplemented with 20 mM MES (Sigma-Aldrich, #M3671) or 20 mM HEPES (Sigma-Aldrich, #H3375, adjusted to the indicated pH values using HCl or NaOH)) for 10 minutes at room temperature. pH values ranged from 5.0 to 7.0 in 0.5-unit increments. Each well received 30 µL of buffer at the indicated pH.

Excitation spectra were collected using a ClarioStar microplate reader (BMG LabTech) with the following parameters: top read, 40 flashes/ well, excitation start: 385 to 10 nm, excitation end: 475 to 10 nm, 15 nm steps; emission: 520 to 10 nm; instrument gain: 2750; focal height 7 mm. Raw fluorescence values at 385 and 475 nm were used to calculate the pHluorin ratio (385/475 nm) for each technical replicate.

### Glosensor Assay for Cellular cAMP Measurement

HEK293T cells were cultured and transiently transfected as described above with 2μg pGloSensor-22F (Promega) and 1μg pcDNA3.1(+) containing the respective A2AR variant. At 24 hours post-transfection, cells were plated in 384-well white clear-bottom plates at a density of approximately 15,000 cells per well in 40 μL of DMEM supplemented with 1% dialyzed FBS (dFBS). Another 18-24 hours later, the culture medium was gently aspirated, and cells were treated with 25 μL per well of the assay buffer solution (Hank’s Buffered Saline Solution; Gibco, 20mM HEPES pH 7.4) containing desired concentration of NECA (Cayman Chemical) supplemented with 1.0 IU/mL adenosine deaminase (Roche) and 2 mM luciferin (GoldBio). Buffer pH was adjusted using KOH at room temperature. Luminescence (no filter) units were collected using the ClarioStar with gain: 3600; focal height 3.5mm. Luminescence data were normalized to percent maximal response and fold change over basal. Dose-response curves were generated by pooling data and fitting to a four-parameter logistic function using GraphPad Prism (version 10.4.0).

## Author contributions

D.G.I. managed the study, and D.G.I. and K.D.L. wrote the manuscript. K.D.L., S.T., A.W. and B.G. performed all experiments.

## Acknowledgements

This work was supported by the National Institutes of Health through an R35 Maximizing Investigators’ Research Award (R35GM119518 to D.G.I.), which provided core funding for this study. Additional support was provided by a Pap Corps Champions for Cancer Research Endowed Chair to D.G.I. and the Sylvester Comprehensive Cancer Center.

